# Rethinking the Benign Nature of Class II Lupus Nephritis

**DOI:** 10.64898/2026.09.13.751280

**Authors:** Jasmine Shwetar, Andrea Fava, Sarah Keegan, Philip M. Carlucci, Jeffrey N. Dudley, Lodoe Lama, Hemant Suryawanshi, Pavel Morozov, Briana J. Mullins, Abhimanyu Amarnani, Linda Procell, Dania Annuar, Siddarth Gurajala, Peter M. Izmirly, Judith A. James, Joel M. Guthridge, Thomas M. Eisenhaure, Arnon Arazi, Chaim Putterman, Deepak A. Rao, Betty Diamond, H. Michael Belmont, William Apruzzese, Anne Davidson, Soumya Raychaudhuri, Nir Hacohen, Maria Dall’Era, Cindy Loomis, Robert M. Clancy, Thomas Tuschl, Ming Wu, Derek M. Fine, Mohamed G. Atta, Jose Monroy-Trujillo, Richard Furie, Diane L. Kamen, Kenneth Kalunian, Accelerating Medicines Partnership in RA/SLE, Michelle Petri, Jill P. Buyon, Brad H. Rovin, Kelly V. Ruggles

## Abstract

**Introduction:** Class II lupus nephritis (LN) is considered clinically benign, yet up to 50% of patients progress to more severe disease and no molecular characterization exists. We aimed to define the single-cell landscape of Class II LN and identify cellular and transcriptional features associated with kidney outcome.

**Methods:** We performed single-cell RNA sequencing on kidney biopsies from 15 Class II LN patients (8 de novo, 7 regressed from prior proliferative or membranous disease) and 6 healthy controls, integrated with 155 LN and 30 healthy-control samples from the Accelerating Medicines Partnership in SLE spanning proliferative, membranous, and mixed LN. Findings were correlated with 52-week renal response and supported by urinary proteomics.

**Results:** Despite histologically minimal changes, Class II LN exhibited marked immune cell expansion comparable to proliferative and membranous LN, including CCL3+ CCL4+ intermediate monocytes producing TNF, FOLR2+ SIGLEC1+ macrophages producing TGF-beta and PDGF, and CD69+ CD8+ effector memory T cells producing IFN-gamma. Fibroblasts and myofibroblasts were significantly expanded and displayed divergent transcriptional programs: pro-fibrotic and inflammatory myofibroblast modules were associated with worse 52-week outcomes, while interferon-responsive and matrix-remodeling fibroblast programs were associated with improvement. Pathogenic and protective fibroblast populations received overlapping immune-derived signals, suggesting that fibroblast transcriptional state, rather than the identity of incoming signals, shapes the stromal response. Baseline chronicity index captured this stromal heterogeneity and was the clinical parameter most associated with outcome. Urinary proteomics aligned Class II with proliferative rather than membranous disease.

**Conclusions:** Class II LN harbors substantial molecular activity not captured by routine histology. Fibroblast transcriptional programs and baseline chronicity index are candidate correlates of kidney outcome that warrant validation in larger, prospectively followed cohorts.

**Lay summary:** Class II lupus nephritis is generally regarded as benign and is not specifically treated, yet up to half of patients later progress to more severe nephritis. To understand this discordance, we applied single-cell RNA sequencing to kidney biopsies from 15 patients with Class II disease and compared them with the full spectrum of lupus nephritis. Despite their mild histologic appearance, Class II kidneys showed immune infiltration comparable to proliferative disease, together with expanded fibroblast populations whose transcriptional programs differed between patients. These fibroblast programs, and the degree of chronic damage already present at biopsy, tracked with kidney outcomes at one year more closely than standard clinical measures. These findings indicate that Class II lupus nephritis carries greater molecular activity than its histology suggests, and that chronicity warrants closer attention in its assessment.

## Introduction

Systemic lupus erythematosus (SLE) is a chronic autoimmune disorder characterized by autoantibody production, lymphoproliferation, and immune complex deposition, resulting in multiorgan inflammation and damage^1^. Among the various SLE manifestations, lupus nephritis (LN) is common, with an incidence of 40-60%^2,3^. LN sharply increases the morbidity and mortality of lupus and may result in kidney failure requiring dialysis or transplantation. However, even with aggressive treatment and the approval of belimumab, voclosporin, and recently obinutuzumab as targeted therapeutics, less than half of patients achieve a complete renal response and 15% develop end-stage kidney disease (ESKD)^4–6^.

The International Society of Nephrology (ISN) and Renal Pathology Society (RPS) classify LN into six distinct classes based on histopathological changes in the glomeruli^7^. Class II LN has been traditionally considered a less aggressive pathology with a favorable prognosis. Based on ISN/RPS criteria, Class II lacks the endocapillary hypercellularity and subendothelial immune complexes of proliferative LN (classes III, IV) and the basement membrane thickening and subepithelial deposits of membranous LN (class V). Due to its perceived benignity, most society guidelines do not suggest specific treatment recommendations for Class II LN and management is generally conservative, favoring a watchful rather than actionable approach. Although Class II LN may resolve or remain stable without aggressive or even modest immunosuppression, several longitudinal studies have shown that up to 50% of patients initially diagnosed with Class II eventually manifest a more severe histologic phenotype^8–11^. This raises the question of whether histology fully captures the biology of Class II LN. We therefore hypothesized that routine histology underestimates the molecular activity present in Class II LN, and set out to define its single-cell landscape and to identify cellular and transcriptional features associated with 52-week kidney outcome.

To address this, we performed high-resolution single-cell RNA sequencing (scRNA-seq) of kidney biopsies with matched longitudinal clinical data from Class II patients, 8 with de novo Class II LN and 7 with Class II LN following a previous diagnosis of proliferative or membranous LN. Additionally, we leveraged the Accelerating Medicines Partnership (AMP) in SLE, which includes transcriptomic data from proliferative and membranous LN kidney biopsies, to characterize the molecular and cellular phenotypes of Class II LN in the context of the full range of LN histopathology^12^.

## Methods

The study was approved by the NYU Institutional Review Board, and written informed consent was obtained from all subjects. Kidney biopsies from 15 Class II LN patients and 6 healthy controls were processed for single-cell RNA sequencing and integrated with 155 LN and 30 healthy-control samples from the Accelerating Medicines Partnership in SLE^13^. Detailed methods are provided in Supplementary Methods. Patients or the public were not involved in the design, conduct, reporting, or dissemination of this research.

## Results

### Clinical characterization and single-cell transcriptomic profiling of Class II Lupus Nephritis

To evaluate the cellular and molecular changes of Class II LN, kidney needle biopsy cores were collected from 15 Class II patients for scRNA-seq. Histologic characteristics of the biopsies and the clinical and demographic characteristics of the patients are shown in Table 1. These 15 Class II LN biopsies were categorized into two groups: 8 patients were classified as ‘*de novo*’ cases, which included those with either a first-time LN diagnosis or stable Class I/II LN from a previous biopsy. The remaining 7 patients were classified as ‘regressed’ cases, representing patients whose disease had improved to Class II following treatment of previously diagnosed proliferative, membranous, or mixed LN (**Fig. 1A and Table 1**). Patients were also evaluated based on their kidney status 52 weeks post-biopsy. Of the 15 patients, nine met criteria for improvement at week 52 (defined, following the ACCESS trial, as UPCR < 0.5 together with serum creatinine that was normal (≤ 1.3 mg/dL) or, if abnormal, ≤ 125% of baseline), while 6 did not, forming two outcome groups: ‘improved’ (n=9) and ‘stable/worse’ (n=6). Notably, clinical findings at baseline such as serum creatinine, estimated glomerular filtration rate (eGFR) and urine protein creatinine ratio (UPCR) had a large range across Class II patients, with two de novo patients having nephrotic range proteinuria (UPCR > 3) (**Fig. 1B**). Class II sample biopsies were evaluated for chronicity index (CI) for features of chronic damage and grouped into none (N=2), mild (N=8), moderate (N=5), and severe (N=0) CI (methods, **Fig. 1B**). De novo patients predominantly had none or mild CI (7/8 patients) while regressed patients showed more moderate CI (4/7 patients), though this difference was not statistically significant (Fisher’s exact test, p = 0.2). Of all clinical parameters analyzed, chronicity at the time of biopsy emerged as the only variable associated with response status at 52 weeks, trending toward significance (p = 0.089). All other clinical parameters at baseline, including previous biopsy status, eGFR, and UPCR showed no significant associations with kidney status at one year.

**Figure 1.**
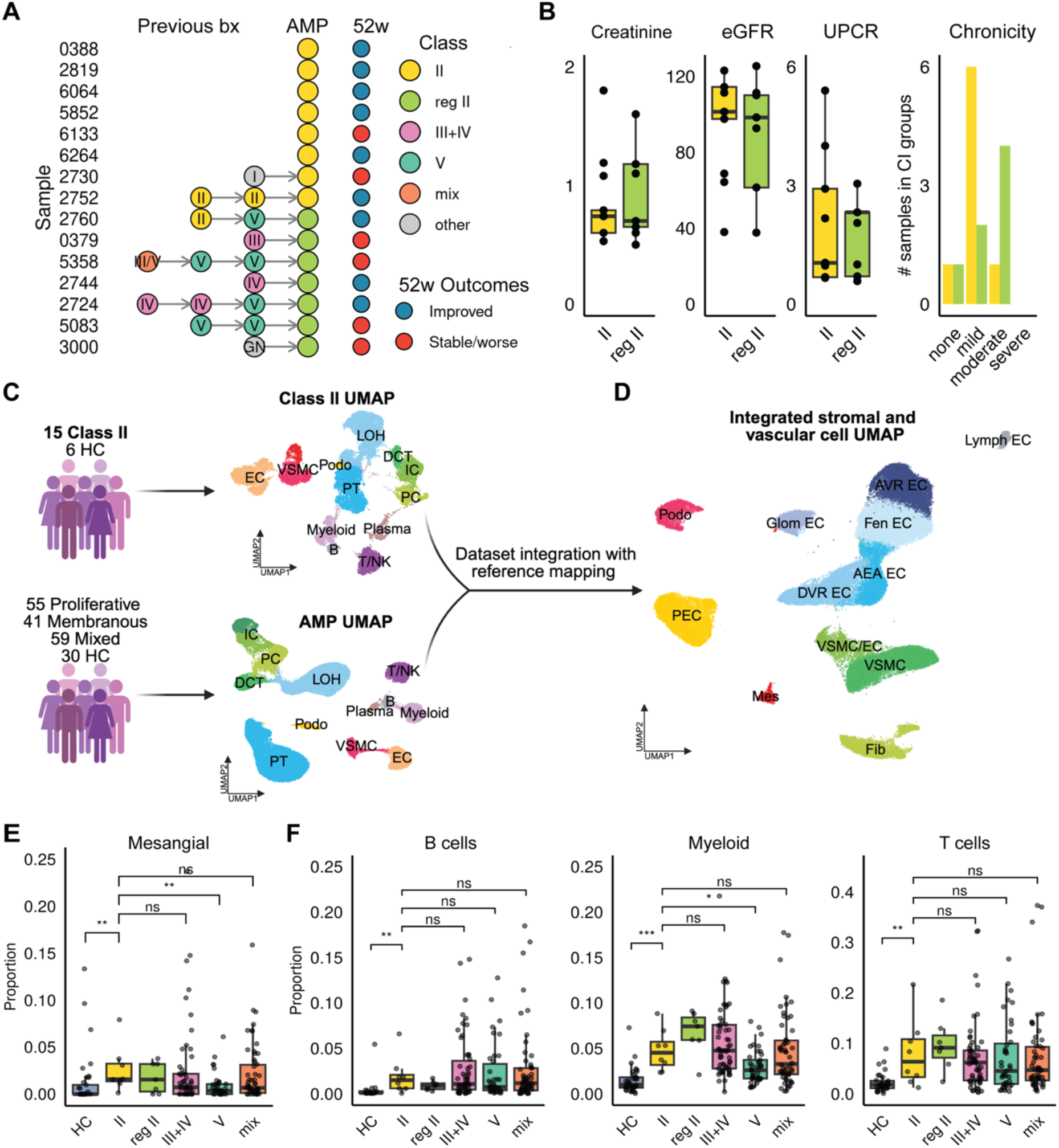
Clinical characterization and single-cell transcriptomic profiling of Class II Lupus Nephritis. (**A**) Class II sample disease history classification and clinical outcomes. Patients were categorized as de novo (yellow; n=8; first-time LN diagnosis or stable Class I/II from previous biopsy) or regressed (green; n=7; improved to Class II following treatment of previously diagnosed proliferative, membranous, or mixed LN). 52-week post-biopsy kidney outcomes are indicated as improved or stable/worse. (**B**) Baseline clinical parameters for Class II patients: serum creatinine (mg/dL), estimated glomerular filtration rate (eGFR; mL/min/1.73 m²), urine protein-to-creatinine ratio (UPCR; mg/mg), and NIH Chronicity Index (CI) categorized as none, mild, or moderate. (**C**) Dataset integration strategy. Class II kidney biopsies (n=15) and AMP datasets were individually clustered, then integrated during subclustering using a reference-based mapping approach. Right panels show total cell UMAPs for Class II (top; 15 Class II and 6 HC samples) and AMP (bottom; 55 proliferative, 41 membranous, 59 mixed LN, and 30 HC samples). Twelve distinct cell populations were identified: proximal tubule (PT), loop of Henle (LOH), distal convoluted tubule (DCT), principal cells (PC), intercalated cells (IC), endothelial cells (EC), vascular smooth muscle cells (VSMC), fibroblasts (Fib), podocytes (Podo), B cells, T cells, and myeloid cells. (**D**) UMAP visualization of integrated stromal and glomerular cell subclustering, identifying 5 endothelial cell types (Ascending and Descending Vasa Recta (AVR/DVR EC), Fenestrated (Fen EC), Afferent/Efferent Arteriole (AEA EC), Glomerular (Glom EC) and Lymphatic (Lymph EC)), Parietal Epithelial Cells (PEC), Podocytes (Podo), Mesangial (Mes), Fibroblasts (Fib), Vascular Smooth Muscle Cells (VSMC) and Endothelial-like VSMCs (VSMC/EC). (**E**) Mesangial cell proportions across disease states, normalized relative to all stromal and glomerular cell types in (D). (**F**) Immune cell proportions (B cells, T cells, myeloid cells) across disease states.

**Table 1.** Demographic Data for Class II samples.

| Sample | Age | Sex | Race | Ethnicity | Chronicity group | IFTA | GGS | Random UPCR | Serum Creatinine | 52 week UPCR | 52 week Creatinine | 52 weeks | Previous bx status | Chronicity group |
| --- | --- | --- | --- | --- | --- | --- | --- | --- | --- | --- | --- | --- | --- | --- |
| 388 | 67 | F | W | NH | none | 0 | 6 | 0.97 | 0.53 | UD | 0.64 | improved | de novo | none |
| 2819 | 35 | F | B | NH | mild | 10 | 10 | 0.66 | 1.08 | 0.042 | 0.9* | improved | de novo | mild |
| 6064 | 47 | F | A | NH | mild | 5 | 2 | 5.4 | 0.6 | 0.09 | 0.56 | improved | de novo | mild |
| 5852 | 39 | F | W | H | mild | 25 | 11 | 2.2 | 0.79 | 0.23 | 0.76 | improved | de novo | mild |
| 6133 | 26 | F | Other | NH | mild | 10 | 0 | 2.9 | 0.74 | 2.2 | 2.9 | stable/worse | de novo | mild |
| 6264 | 27 | F | B | NH | mild | 7.5 | 7 | 1 | 1.2 | neg UA** | 0.92 | improved | de novo | mild |
| 2730 | 32 | F | B | NH | moderate | 50 | 24 | 4 | 1.8 | 1 | 4.2 | stable/worse | I | moderate |
| 2752 | 31 | F | B | NH | mild | 10 | 0 | 0.68 | 0.6 | 0.16 | 0.7 | improved | II;II | mild |
| 2760 | 35 | F | A | NH | none | 3 | 0 | 1 | 0.5 | 0.36 | 0.5 | improved | II;V | none |
| 379 | 44 | F | B | NH | mild | 25 | 25 | 0.7 | 0.65 | 1.6 | 0.64 | stable/worse | III | mild |
| 5358 | 36 | F | W | NH | moderate | 20 | 45 | 3 | 1.2 | 0.27 | 1.5 | stable/worse | III/V;V;V | moderate |
| 2744 | 67 | F | B | NH | mild | 15 | 19 | 0.58 | 0.6 | 0.31 | 0.7 | improved | IV | moderate |
| 2724 | 30 | M | W | NH | moderate | 25 | 33 | 2.05 | 1.1 | 0.08 | 1.3 | improved | IV/IV/V | moderate |
| 5083 | 43 | F | B | NH | moderate | 20 | 35 | 2.32 | 0.7 | 1.8 | 0.68 | stable/worse | V;V | moderate |
| 3000 | 56 | F | W | H | moderate | 30 | 69 | 0.7 | 1.6 | 0.2 | 2.2 | stable/worse | GN | moderate |

**Legend/Abbreviations:** IFTA, interstitial fibrosis and tubular atrophy; **GGS**, global glomerulosclerosis; UPCR, urine protein-to-creatinine ratio (mg/mg); UD, undetectable; NR, no response; GN, glomerulonephritis; W, White; B, Black; A, Asian; H, Hispanic; NH, non-Hispanic; F, female; M, male.
\*85 week response status
\*\*UPCR not done, negative on Urine Analysis (UA)

To understand the molecular pathology of Class II LN and compare it to other more advanced LN classes, we integrated these data with a complementary AMP dataset comprising 55 proliferative, 41 membranous, 59 mixed, and 30 HC samples (**Fig. 1C**)^13^. Datasets were integrated during sub-clustering analysis using a reference-based mapping approach, and marker expression confirmed the same cell-states were captured across both datasets (**Fig. 1D**; Fig. S1A and B). Collectively, this study comprises 568,547 high-quality cells from 170 LN kidneys across the full LN histopathologic spectrum.

### Class II LN exhibits mesangial and immune expansion

The histologic hallmark of Class II LN is the presence of immune complexes in the glomerular mesangium with mesangial cell proliferation. To normalize for variations in biopsy quality and disease-specific factors such as increased immune infiltration, proportions were analyzed relative to all other cell types in subclustering analysis (**Fig. 1D**; Fig. S1B). Both regressed and de novo Class II LN showed significantly higher mesangial cell proportions compared to HCs (p = 0.0012 de novo vs HC comparison, p = 0.052 regressed vs HC), consistent with histological requirements for a Class II diagnosis (**Fig. 1E**). While Class II mesangial proportions were comparable to proliferative (Class III+IV) and mixed LN, both of which can exhibit varying degrees of mesangial hypercellularity, they were significantly elevated compared to membranous LN (Class V).

Despite Class II’s histological definition of minimal immune cells, single-cell analysis revealed significant immune cell expansion in both de novo and regressed Class II samples (**Fig. 1F**). B cells, T cells, and myeloid cells were all markedly elevated compared to HCs, with abundances comparable to proliferative, membranous, and mixed LN classes. Analysis of absolute cell counts confirmed that these immune cell expansions represent true increases rather than proportional artifacts (Fig. S1D). This substantial immune infiltration revealed active inflammatory processes in Class II LN that are not captured by routine histological assessment.

### Distinct inflammatory and pro-fibrotic myeloid phenotypes characterize Class II LN

Among all immune cell types examined, myeloid cells exhibited the most pronounced changes in Class II kidneys. Sub-clustering analysis identified 17 distinct populations including monocytes (M1–M5), macrophages (M6–M10), and dendritic cells (**Fig. 2A and B**).

**Figure 2.**
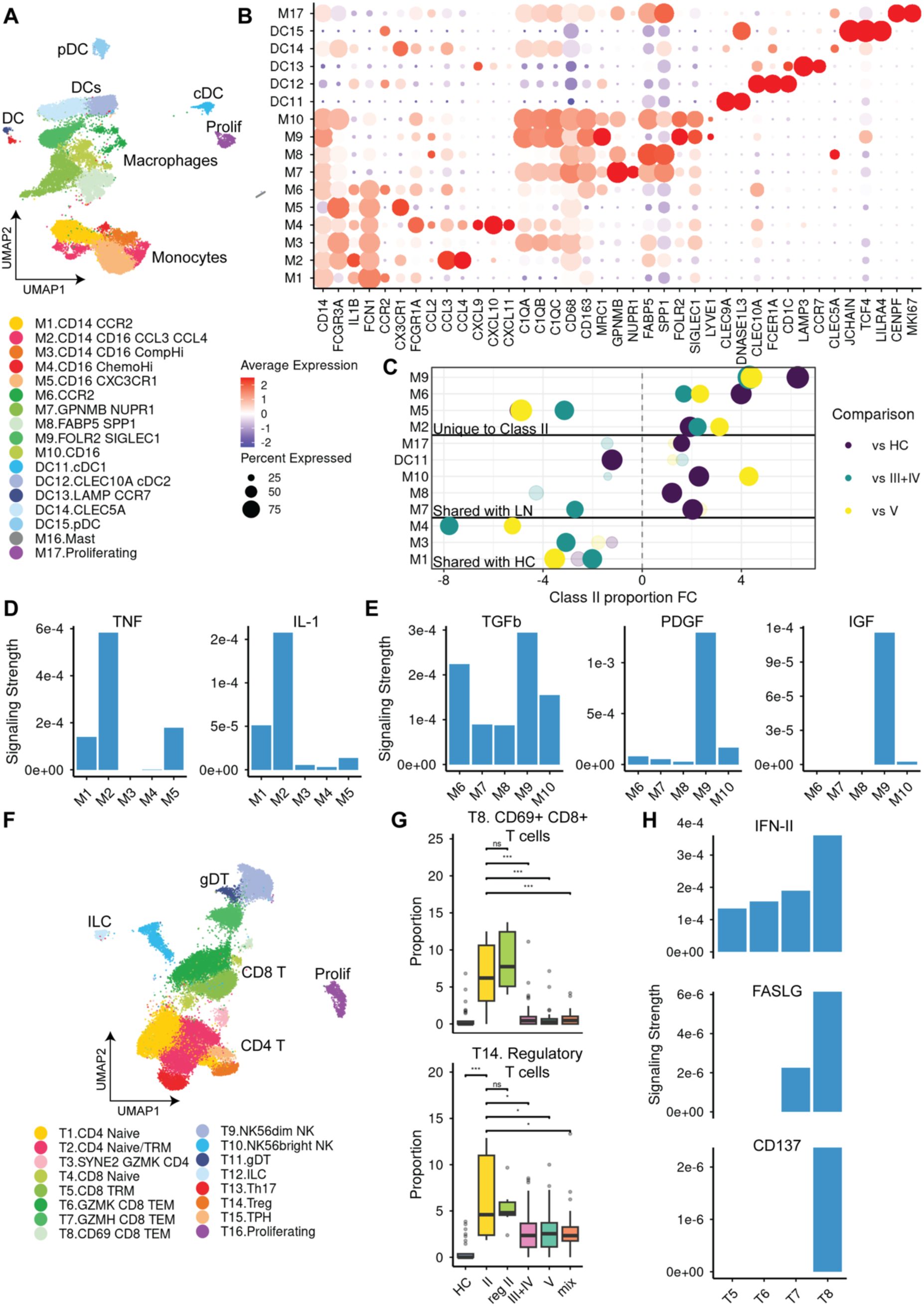
Distinct myeloid and T cell phenotypes characterize Class II Lupus Nephritis. (**A**) Subclustering of myeloid lineage cells from all LN and HC samples, identifying 17 distinct populations including monocytes (M1-M5), macrophages (M6-M10), and dendritic cells. (**B**) Expression of canonical markers defining monocyte and macrophage subsets. (**C**) Myeloid cell type proportional differences across disease states. Fold change (FC) comparing de novo Class II LN to healthy controls (HC), proliferative (Class III+IV), and membranous (Class V) LN. Dot size represents statistical significance (-log10 p-value) from Wilcoxon rank-sum tests, with larger dots indicating stronger significance. Colors distinguish between comparisons, while cell types are grouped based on whether their patterns are not significantly different from HC, shared with proliferative/membranous LN, or unique to Class II. All values calculated relative to de novo Class II; any differences between de novo and regressed immune cells are addressed in the text. Only cell types with statistically significant differences from Class II are shown. (**D**) Ligand signaling strength for TNF and IL1 pathways in monocyte populations (M1-M5). (**E**) Ligand signaling strength for TGF-beta, PDGF, and IGF pathways in macrophage populations (M6-M10). (**F**) Subclustering of T and NK lineage cells, identifying 16 distinct populations. (**G**) Quantification of T8 CD69+ CD8+ effector memory T cells and T14 regulatory T cell proportions across disease states. (**H**) Ligand signaling strength for IFN-II, FASLG, and CD137 pathways in effector CD8+ T cell populations (T5-T8). See also Figure S2.

Compositional analysis revealed distinct myeloid alterations in Class II LN (**Fig. 2C**; Fig. S2A). All monocyte populations were depleted in Class II except for M2 CCL3+ CCL4+ intermediate monocytes, an inflammatory chemokine-expressing subset, which were significantly elevated in both de novo and regressed Class II compared to HCs and all other LN classes. Among macrophage populations, M6 monocyte-like macrophages and M9 FOLR2+ SIGLEC1+ macrophages, a tissue-resident population marked by interferon-responsive genes, were uniquely expanded in both de novo and regressed Class II compared to HCs and all other LN classes (**Fig. 2G**). M10 CD16+ macrophages were the only myeloid subset that differed between de novo and regressed Class II, with de novo showing significantly higher proportions comparable to proliferative and mixed LN, while regressed Class II and membranous LN had significantly fewer of these cells (Fig. S2A). These Class II-enriched populations expressed key inflammatory and fibrotic mediators: M2 intermediate monocytes expressed *TNF* and *IL1B* (**Fig. 2D**), while M9 macrophages expressed *TGFB1*, *PDGFA*, and *PDGFB* (**Fig. 2E**).

### Class II LN shows selective expansion of activated effector and regulatory T cells

To elucidate how T cells and Natural Killer (NK) cells contribute to the immune environment in Class II LN, we performed detailed sub-clustering analysis, revealing 16 distinct states based on canonical markers (**Fig. 2F**; Fig. S2B). Compositional analysis revealed fewer T cell alterations in Class II kidneys compared to the pronounced myeloid changes (Fig. S2C). Two T cell populations showed unique Class II-specific expansion: T8 CD69+ CD8+ effector memory T cells (TEM) and T14 regulatory T cells (Tregs) were both significantly elevated in de novo and regressed Class II compared to HCs and all other LN classes. T8 TEM expressed *IFNG*, *TNFSF9* (encoding 4-1BB ligand), and *FASLG* (encoding Fas ligand) (**Fig. 2H**), indicating active effector T cell responses within Class II kidney tissue.

Collectively, these immune populations expressed key pro-fibrotic and inflammatory mediators (TNF, TGF-beta, PDGF, and IFN-gamma). Although Class II LN is not typically associated with significant interstitial fibrosis, these cytokines are canonical activators of fibroblasts and tissue remodeling, prompting examination of their abundance and transcriptional states.

### Class II LN exhibits significant fibroblast expansion with heterogeneous activation states

To investigate the stromal response to immune-derived signals, we examined fibroblast proportions using the integrated stromal and glomerular cell UMAP (**Fig. 1D**). Both de novo and regressed Class II cases exhibited significantly higher fibroblast proportions compared to HCs and all other LN classes (**Fig. 3A**; p < 0.01), confirmed by absolute cell counts (Fig. S3A). To assess this expansion in intact tissue, we performed immunofluorescence staining using fibroblast-specific protein 1 (FSP1, *S100A4*) and proliferation marker Ki-67. Analysis of two de novo Class II and two proliferative biopsies showed comparable percentages of double-positive S100A4+ Ki-67+ proliferating fibroblasts (**Fig. 3B**). Despite the expansion of fibroblasts, their proportions in Class II LN biopsies showed no correlation with baseline eGFR or UPCR (**Fig. 3C**).

**Figure 3.**
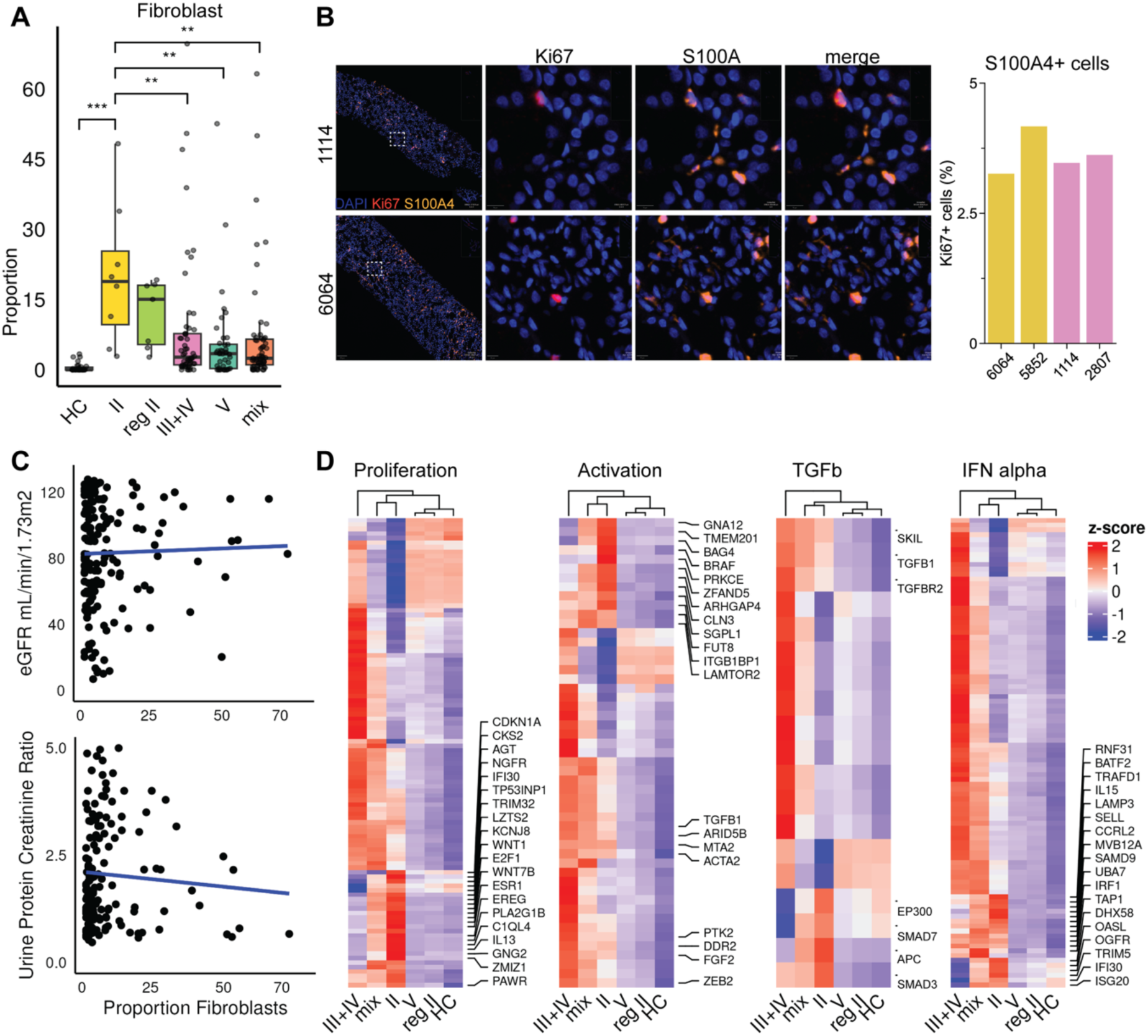
Class II LN exhibits significant fibroblast expansion with heterogeneous activation states. (**A**) Per sample proportion of fibroblasts relative to all cells, across disease states. (**B**) Immunofluorescence staining for Ki-67 and fibroblast-specific protein 1 (FSP1/S100A4) in de novo Class II and proliferative LN biopsies (left) with quantification of double-positive S100A4+ Ki-67+ proliferating fibroblasts (right). Color coding as in (A). (**C**) Linear regression of eGFR and UPCR versus proportion of fibroblasts per sample. (**D**) Per-class aggregated scaled expression in fibroblasts of proliferation/migration, activation, TGF-beta signaling, and IFN-alpha response gene signatures.

To better understand the activation state of fibroblasts in Class II, we analyzed four key gene programs: a combined fibroblast proliferation and migration signature, activation markers, TGF-beta signaling, which drives fibrosis, and IFN-alpha response, a central mediator of lupus pathogenesis^14,15^. Fibroblasts from de novo Class II biopsies showed consistently elevated expression across all four gene programs, with patterns most similar to the high expression observed in mixed (proliferative plus membranous) nephritis (Fig. 3D). In contrast, fibroblasts from regressed Class II biopsies showed expression patterns more similar to membranous LN. These bulk pathway scores average across a heterogeneous fibroblast population; sub-clustering and NMF analysis below resolve this heterogeneity by identifying distinct co-expression modules with divergent clinical associations. This relationship was fibroblast-specific and not observed in other cell types such as mesangial cells (Fig. S3B).

### Fibroblast gene programs identify distinct functional states with clinical associations

To better understand fibroblast heterogeneity in Class II LN, we performed subclustering analysis of fibroblasts with vascular smooth muscle cells (VSMCs), identifying fibroblasts, myofibroblasts (activated fibroblasts with enhanced contractile and matrix-producing capacity) and three VSMC populations (**Fig. 4A**; Fig. S4A). Fibroblasts, defined by high *S100A4* and *DCN*, were expanded in Class II relative to HCs, consistent with earlier observations. In contrast, myofibroblasts showed a Class II-specific expansion pattern, being uniquely elevated in both de novo and regressed Class II compared to both HC and all other LN classes (Fig. S4B and C).

**Figure 4.**
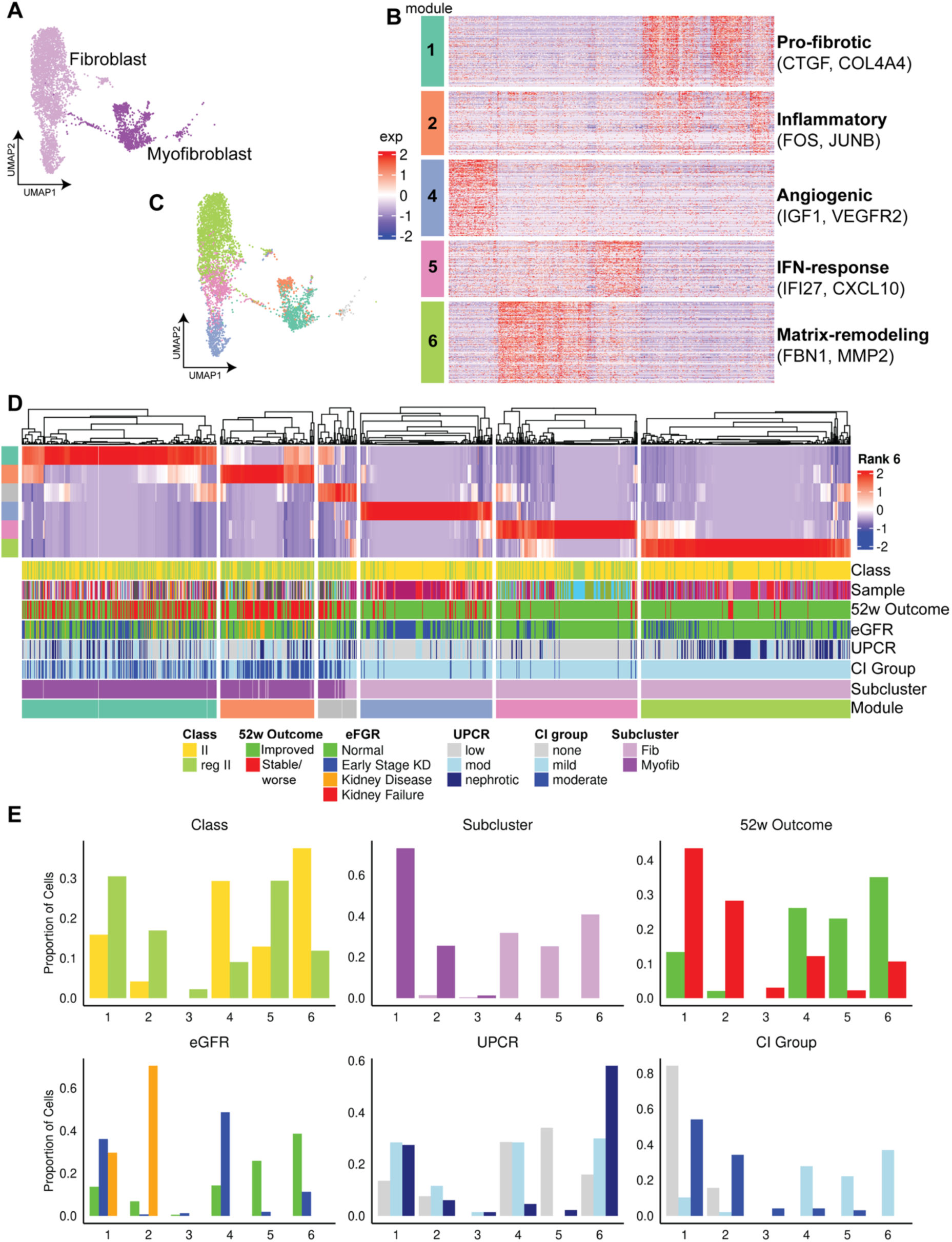
Fibroblast gene programs identify distinct functional states with clinical associations. (**A**) Subclustering of fibroblasts and myofibroblasts (Myofib), identifying fibroblasts, myofibroblasts, and three VSMC populations. (**B**) Scaled expression of genes in 5 NMF gene modules in all fibroblast and myofibroblast cells from Class II samples: Myofib1 (Pro-fibrotic), Myofib2 (Inflammatory), Fib4 (Growth Factor-enriched), Fib5 (Interferon-responsive), and Fib6 (Matrix-remodeling). (Module 3, a mitochondrial technical artifact, excluded). (**C**) Module expression in UMAP space. (**D**) Heatmap of NMF coefficient matrix showing module expression weights across individual Class II fibroblast and myofibroblast cells. (**E**) Proportion of cells from samples within each clinical category expressing each NMF module. Clinical parameters include CI Group (mild/moderate), Class (de novo II/regressed II), eGFR (normal/early-stage kidney disease), 52-week outcomes (Improved/’Stable/Worse’), and UPCR (low/moderate/nephrotic). See also Table 2.

**Table 2.** NMF module co-expression programs and functional characterization.

| Module | Most Representative Genes | Co-expression Program Summary | Functional State |
| --- | --- | --- | --- |
| **Myofib1** | C7, A2M, ITGA1, DACH1, ECRG4, ATRNL1, COL4A4, THSD4 | TGFβ/PDGF signaling receptors (PDGFRA, NRP1, EDNRB)<br>Pro-fibrotic regulators (CCN2/CTGF, EPAS1/HIF2α)<br>Structural collagens (COL4A3/4, COL18A1, COL27A1, COL23A1, COL25A1)<br>Matrix remodeling enzymes (ADAMTS9/12/15)<br>Angiogenic factors (ANGPT1, PTN, HGF, EPO)<br>Inflammatory signaling (STAT4) | Pro-fibrotic myofibroblast program integrating growth factor response with ECM production |
| **Myofib2** | C11orf96, JUNB, APOE, FOS, HSPA1A, GEM | AP-1 transcription complex (FOS/FOSB/JUN/JUNB)<br>Inflammatory transcription networks (RELB/REL, NFATC1/2, IRF1/8, SOX4)<br>TNF pathway components (TNFRSF12A, TNFAIP3/6, TRAF1)<br>Heat shock proteins (HSPA1A/B, HSPB1, HSP90AA1)<br>Broad-spectrum chemokines (CCL2/5/8/13/19/21, CXCL2/8) | Inflammatory myofibroblast program with stress response and immune recruitment |
| **Fib4** | PDZRN4, ADIRF, SYT1, RSPO3, RARRES1, COL8A1, HPGD, SHISA3, FIBIN, | Growth factor signaling (IGF1/2, KITLG, BMP3)<br>Angiogenic receptors (KDR/VEGFR2, NRP2)<br>Wnt potentiators (RSPO3/1)<br>Stress response genes (NOX4, | Growth factor-enriched program with stress response and angiogenic signaling |
|  | ADAMTS3, CD9, EFNA5, MAP3K5, IGF1, OMD | DDIT4, TXNIP, MAP3K5/ASK1)<br>Developmental guidance cues (EFNA5, SEMA6D) |  |
| **Fib5** | SFRP2, CCDC80, IFI27, APOD, SRPX | Interferon-stimulated genes (IFI27, IFI44L, IFIT1/3, MX2, OAS1, BST2, CXCL10)<br>Immune-organizing factors (TNFSF13B/BAFF, VCAM1, CXCL12/14)<br>Wnt inhibitors (SFRP2, DKK2)<br>Anti-inflammatory mediators (SERPINF1, CLU)<br>Metabolic regulators (PPARG, FABP4) | Interferon-responsive program with immune-organizing and anti-inflammatory features |
| **Fib6** | MFAP5, FBN1, SEMA3C, S100A4, SCARA5, EBF2, CREB5, S100A10, SDK1, CNTN4, CD248, GAS7, SLPI | Structural ECM components (FBN1, MFAP5, TNXB, FN1, NID1)<br>Matrix proteases (MMP2/16, ADAMTS5/6/16)<br>S100 calcium-binding proteins (S100A4/6/10/11/13/16)<br>Immune signaling molecules (C3, CSF1, IL18, CD70, TNFSF9)<br>Activated fibroblast markers (CD248, CD34, PDPN) | Matrix-organizing program with active remodeling and immune signaling |

To further characterize functional heterogeneity at higher resolution, we performed Non-negative Matrix Factorization (NMF), which identifies coordinated gene co-expression patterns without imposing artificial boundaries between cell states^16^. This revealed 6 distinct gene expression modules (**Fig. 4B and Table 2**), with Module 3 (mitochondrial genes) excluded as a technical artifact. Module expression patterns in UMAP space showed Modules 1-2 (Myofib1, Myofib2) localizing to myofibroblasts and Modules 4-6 (Fib4, Fib5, Fib6) enriched in fibroblasts (**Fig. 4C**).

The myofibroblast-associated modules represented two distinct pathogenic programs: Myofib1 (Pro-fibrotic) coordinated TGF-beta/PDGF receptor signaling with collagen production and ECM remodeling, while Myofib2 (Inflammatory) coordinated AP-1 transcription factor activation with TNF pathway signaling and chemokine production. The fibroblast modules represented three alternative functional states: Fib4 (Growth Factor-enriched), Fib5 (Interferon-responsive), and Fib6 (Matrix-remodeling) (Table 2).

To investigate whether fibroblast functional states associated with clinical outcomes, we analyzed NMF module expression patterns relative to baseline clinical measurements and 52-week kidney status (**Fig. 4D and E**). Pathogenic myofibroblast modules were enriched in regressed Class II (Myofib1 OR=2.19; Myofib2 OR=4.49), in moderate versus mild chronicity (Myofib1 OR=10.13; Myofib2 OR=10.52), and in patients with stable/worse versus improved 52-week kidney status (Myofib1 OR=4.64; Myofib2 OR=8.10). Protective fibroblast modules showed the opposite pattern. Fib6 was enriched in de novo Class II (OR=4.84), absent in moderate chronicity, and associated with improved outcomes (OR=6.32). Fib5 was modestly enriched in regressed Class II (OR=1.93) but depleted in moderate chronicity (OR=0.10) and strongly associated with improved outcomes (OR=11.11). Fib4 was enriched in de novo Class II (OR=2.54) and early-stage kidney disease (OR=3.99) but depleted in nephrotic-range proteinuria (OR=0.18); its outcome association was weaker (OR=1.68). These associations were fibroblast-specific and not observed in VSMCs (Fig. S4D).

### Immune-stromal signaling networks link specific immune populations to fibroblast functional states

To map potential immune-stromal interactions, we performed ligand-receptor analysis using CellChat to infer cell-cell communication networks^17^. This analysis identified predicted signaling relationships between Class II immune populations and specific fibroblast states (**Fig. 5A and 5B**). Pro-fibrotic Myofib1 and Matrix-remodeling Fib6 were the strongest predicted receivers of TNF signaling from M2 CCL3+ CCL4+ intermediate monocytes and TGF-beta, PDGF, and IGF signaling from M9 FOLR2+ SIGLEC1+ resident macrophages, with Myofib1 as the stronger inferred receiver of TGF-beta, PDGF, and IGF. T8 CD69+ CD8+ effector memory T cells were identified as a predicted source of IFN-gamma and FASLG signaling to both Interferon-responsive Fib5 and Matrix-remodeling Fib6, with Fib6 showing the strongest inferred IFN-gamma reception and Fib5 the strongest FASLG reception.

**Figure 5.**
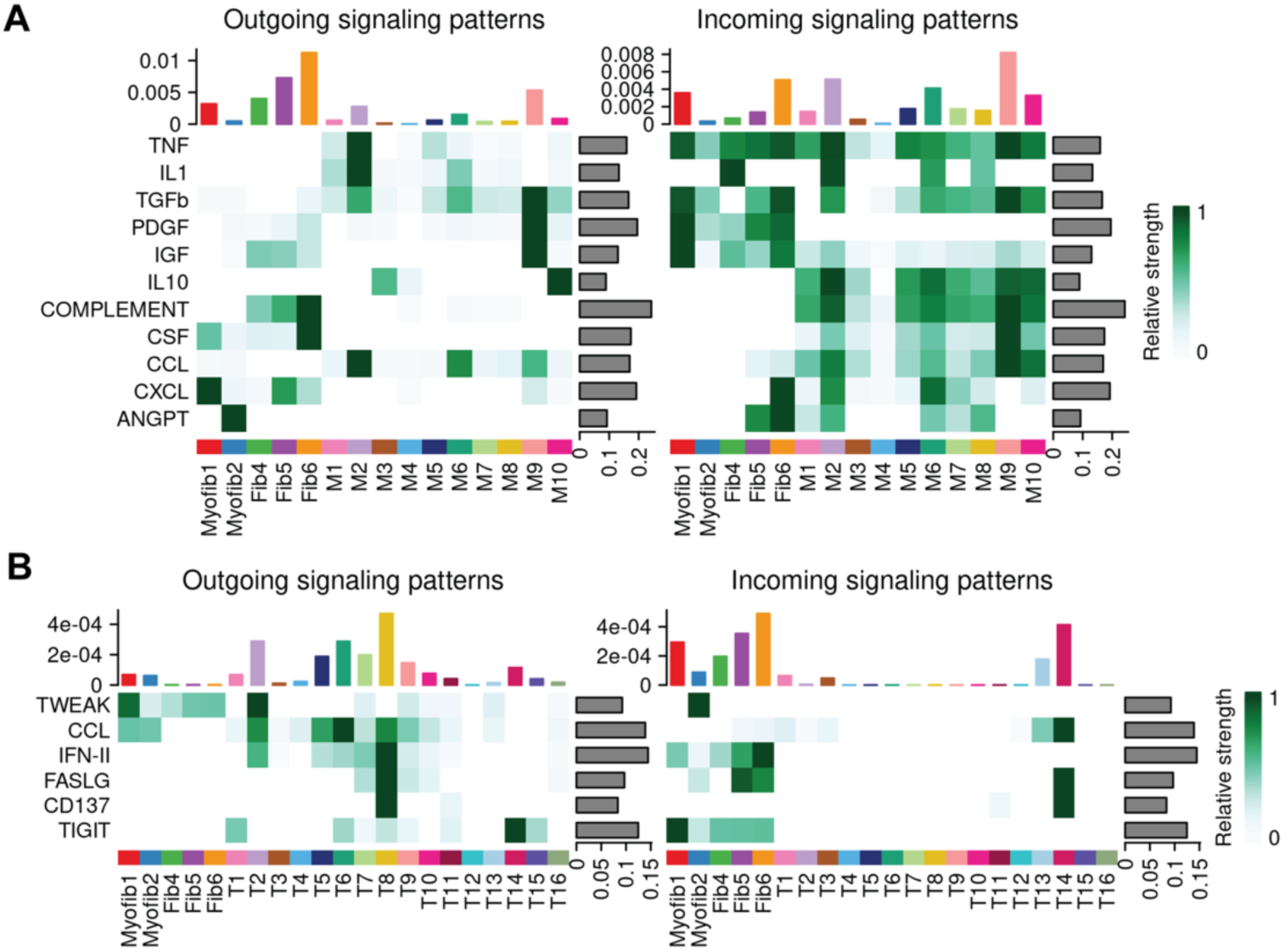
Immune-stromal signaling networks link specific immune populations to fibroblast functional states. (**A**) CellChat-inferred signaling patterns between stromal (Myofib1, Myofib2, Fib4, Fib5, Fib6) and myeloid (M1-M10) populations. Left: outgoing signaling strength. Right: incoming signaling strength. Heatmap shows relative signaling strength for each pathway (TNF, IL1, TGF-beta, PDGF, IGF, IL10, COMPLEMENT, CSF, CCL, CXCL, ANGPT). Bar plots (top) show total signaling strength per cell population; bar plots (right) show total signaling strength per pathway. (**B**) CellChat-inferred signaling patterns between stromal (Myofib1, Myofib2, Fib4, Fib5, Fib6) and T cell (T1-T16) populations. Left: outgoing signaling strength. Right: incoming signaling strength. Pathways shown include TWEAK, CCL, IFN-II, FASLG, CD137, and TIGIT.

### Urinary proteomics recapitulate macrophage and stromal signatures

Given that urine collects the byproducts of renal inflammation and tissue remodeling, we examined whether our single-cell findings were reflected in urine proteomics from Class II LN patients. Proteomic analysis revealed significant upregulation of both fibrotic and immune-related proteins in Class II LN urine compared to HCs. Notably, protein markers characteristic of the expanded M9 FOLR2+ SIGLEC1+ macrophage population were elevated, including CD206 (MRC1), FOLR2, SIGLEC1, LYVE1, and CD163 (**Fig. 6A**).

**Figure 6.**
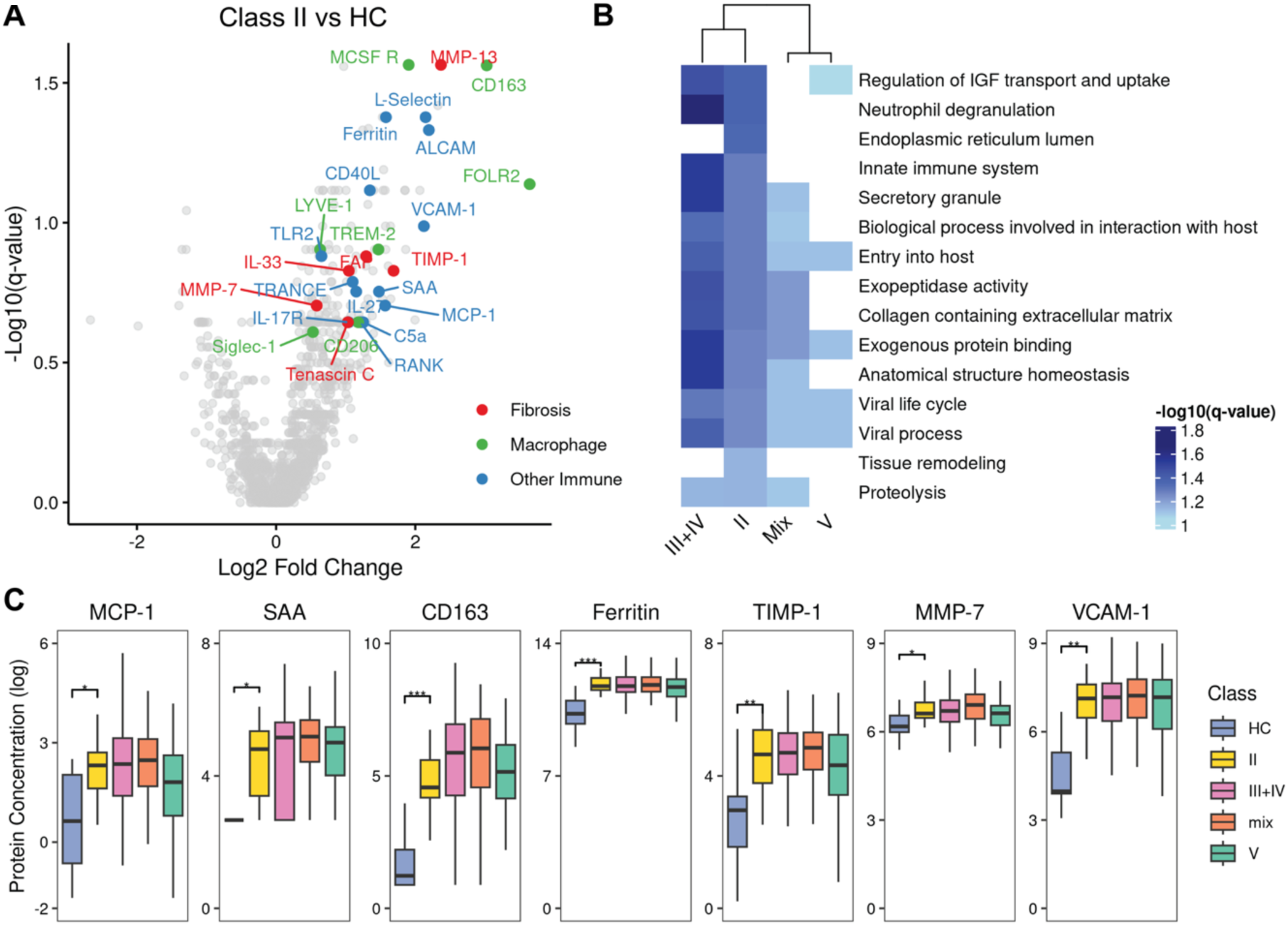
Urine proteomics shows fibrotic and immune signatures. (**A**) Volcano plot of differentially abundant proteins comparing Class II LN urine samples to HCs. Proteins are color-coded by functional category: fibrosis-related proteins (red), macrophage-associated proteins (green), and other immune-related proteins (blue). (**B**) Pathway enrichment analysis of differentially abundant proteins in Class II LN versus HC. Bars represent pathways in Class II, colored by -log10(p-value). Pathways filtered by q-value < 0.1 with top 15 selected by adjusted p-value significance. Significance of the same pathways in Proliferative vs HC, Membranous vs HC, and Mixed vs HC are also shown for comparison. (**C**) Protein concentrations (log-transformed) across LN classes. Significance determined via Wilcoxon rank-sum tests where *p < 0.05, **p < 0.01, ***p < 0.001, ns = not significant.

Pathway analysis of differentially abundant urinary proteins in Class II revealed two major enrichment patterns that paralleled our tissue transcriptional findings (**Fig. 6B**). The first category reflected tissue remodeling processes observed in the stromal compartment, including tissue remodeling pathways, anatomical structure homeostasis, collagen-containing extracellular matrix, proteolysis, and exopeptidase activity. The second category was consistent with the immune activation seen in tissue, showing enrichment of innate immune responses, neutrophil degranulation, and secretory granule function. To determine whether the molecular signature of Class II LN more closely resembled proliferative or membranous disease, we compared urinary protein pathway enrichments across LN classes. The urinary protein signature in Class II LN most closely resembled that of proliferative LN, which also showed enrichment for immune-related processes including neutrophil degranulation, innate immune system, and secretory granule function. In contrast, membranous LN did not demonstrate these immune-related enrichments. Additionally, traditional biomarkers associated with LN and kidney disease (CD163, Ferritin, and TIMP1)^18–20^ were elevated in both Class II and classes III, IV, and V LN compared to healthy controls (**Fig. 6C**).

## Discussion

Class II lupus nephritis lacks specific treatment recommendations across major society guidelines, including EULAR/ERA-EDTA^21^ and KDIGO^22^, reflecting the long-held assumption that mesangial proliferative disease is clinically benign. Yet up to 50% of Class II patients progress to proliferative or membranous LN^8–11^. Through the first single-cell transcriptomic characterization of Class II LN, we find that this histologic class harbors far greater molecular activity than its designation implies: a distinctive immune signature shared across all Class II patients, alongside divergent fibroblast transcriptional programs that were associated with 52-week kidney outcomes. Notably, pathogenic and protective fibroblast populations received overlapping immune-derived signals, pointing to the intrinsic transcriptional state of the fibroblast, rather than the inflammatory input it receives, as the feature that tracked with clinical trajectory.

### Discordance between histology and molecular phenotype

Despite Class II’s histologic designation of minimal immune infiltration, we observed marked B cell, T cell, and myeloid expansion comparable to proliferative and membranous LN (**Fig. 1F**), with these populations producing TNF, TGF-beta, PDGF, and IFN-gamma (**Fig. 2D, E and H**), cytokines with established roles in driving renal inflammation and fibrosis^23,24^. We also observed substantial fibroblast and myofibroblast expansion in Class II (**Fig. 3A and B**; Fig. S4B and C), a finding that was unexpected given the minimal interstitial fibrosis typically seen on light microscopy. Fibroblast proportions did not correlate with baseline eGFR or UPCR (**Fig. 3C**), despite interstitial fibrosis being a well-established predictor of kidney function decline in LN^25–27^. This dissociation likely reflects cohort heterogeneity: some patients presented with moderate chronicity and established fibrosis, while others showed transcriptional fibroblast activation in the absence of significant chronic damage. In the latter group, this pattern is consistent with a pre-fibrotic state: activation without measurable collagen deposition^28,29^, extending the concept of “molecular scars” described in acute kidney injury^30^. This suggests that a subset of Class II patients have ongoing stromal activation that precedes the establishment of measurable fibrosis.

### Fibroblast transcriptional programs are associated with clinical outcomes

All Class II patients shared a core immune signature consisting of M2 CCL3+ CCL4+ intermediate monocytes, M6 monocyte-like macrophages, M9 FOLR2+ SIGLEC1+ macrophages, T8 CD69+ CD8+ effector memory T cells, and T14 regulatory T cells that together distinguished Class II from all other LN classes (**Fig. 2C and G**). The simultaneous expansion of effector T cells and Tregs is consistent with active immune engagement accompanied by counter-regulatory responses, a pattern seen in other autoimmune tissue microenvironments where regulatory and effector populations co-expand during active disease.

Despite receiving these shared immune signals, fibroblasts responded with divergent transcriptional programs, and it was these fibroblast programs, not standard clinical parameters including eGFR and UPCR, that showed the strongest associations with 52-week kidney outcomes. On the pathogenic side, Myofib1 (Pro-fibrotic) represents a canonical TGF-beta/PDGF-driven fibrosis program with collagen production and ECM remodeling, similar to myofibroblast differentiation trajectories identified in human kidney fibrosis^29^. Myofib2 (Inflammatory) showed an even stronger association with worse outcomes (OR=8.10). Its defining feature is expression of TNFRSF12A, encoding the TWEAK receptor Fn14. The TWEAK/Fn14 axis has been directly implicated in LN pathogenesis: TWEAK drives myofibroblast differentiation and fibrogenesis in kidney disease^31^ and urinary TWEAK levels correlate with LN disease activity^32^. Ligand-receptor analysis identified TWEAK as Myofib2’s dominant predicted incoming signal (**Fig. 5B**), positioning TWEAK/Fn14 as the principal predicted route through which the highest-risk module may be engaged.

Among the protective modules, Fib5 (Interferon-responsive) showed the strongest association with improved outcomes (OR=11.11). Notably, IFN-gamma, produced by the expanded T8 effector memory T cells, is also a well-characterized inhibitor of fibroblast-to-myofibroblast differentiation and collagen synthesis^33,34^. Predicted IFN-gamma delivery from T8 cells to both protective fibroblast populations (**Fig. 5B**) points to a role for this cytokine in maintaining non-fibrogenic stromal states. Fib6 (Matrix-remodeling) pairs ECM production with matrix-degrading proteases, resembling resolution-phase fibroblasts in reversible kidney injury^29,30^. Ligand-receptor analysis predicts that Fib6 receives many of the same M9-derived TGF-beta, PDGF, and IGF signals as Myofib1, yet associates with improved rather than worse outcomes. Fibroblast state, then, rather than the identity of incoming signals, appears to shape the functional response.

### Chronicity index as a clinical correlate of fibroblast transcriptional state

The fibroblast programs described above were strongly associated with baseline chronicity index. While pathologists typically note individual features of chronic damage such as tubular atrophy, interstitial fibrosis, and glomerulosclerosis in Class II biopsies, CI is not routinely scored as a composite measure or used to guide management decisions in this class. Our deliberate CI assessment revealed that 5 of 15 Class II patients had moderate chronicity despite a Class II designation, and CI was the only clinical parameter to show an association with 52-week outcomes, though this did not reach statistical significance (p=0.089). Disease history partially overlapped with CI, as regressed patients more commonly presented with moderate CI, although this was not statistically significant. However, Fib5 was modestly enriched in regressed Class II despite being strongly depleted in moderate chronicity. This pattern suggests that molecular state at biopsy, rather than class designation or disease history alone, more closely tracks clinical trajectory.

### A window of opportunity in de novo Class II LN

De novo Class II patients exhibited a molecular signature more reminiscent of proliferative LN than expected from a histologically mild disease. Multiple independent lines of evidence converge to support this: (i) fibroblast activation signatures elevated to levels seen in mixed nephritis (**Fig. 3D**), (ii) urinary proteomic enrichment for immune activation pathways characteristic of proliferative rather than membranous disease (**Fig. 6A and B**), and (iii) mesangial expansion shared with proliferative LN (**Fig. 1E**). This convergence provides a molecular rationale for the clinical observation that up to 50% of Class II patients progress to proliferative disease^8–11^. Despite this proliferative-like profile, de novo patients predominantly presented with none or mild CI and protective fibroblast programs, and were more likely to improve at 52 weeks. De novo Class II may therefore represent a window in which the inflammatory inputs are active but the stromal response remains dominated by protective programs. Whether this protective stromal balance can be preserved, and whether its loss precedes histologic progression, warrants prospective investigation.

### Limitations and future directions

Several limitations should be acknowledged. While our single-cell analysis provides unprecedented resolution of the cellular landscape in Class II LN, the sample size is modest. The association between CI and 52-week outcomes trended toward but did not reach statistical significance, likely reflecting limited statistical power. The cross-sectional design prevented determination of whether fibroblast modules independently predict outcomes or whether CI is the primary driver with modules as its molecular correlate. Treatment was not standardized and effects on fibroblast programs could not be assessed. Our cohort also combined de novo Class II with kidneys that had regressed to Class II following prior proliferative or membranous disease; stromal and chronicity-associated signals in the regressed group may therefore partly reflect prior injury and treatment rather than intrinsic Class II biology, and we reported the two groups separately where they diverged. Longitudinal studies with serial biopsies would establish whether transcriptional programs precede histologic transformation and whether incorporating chronicity assessment refines risk stratification over time. Spatial transcriptomics would directly test the physical proximity of immune and stromal cells inferred from ligand-receptor analysis^35^. Validation in larger, prospectively enrolled cohorts with standardized treatment protocols is needed to confirm whether incorporating chronicity assessment refines risk stratification in Class II LN.

### Conclusions

Our findings challenge the view of Class II LN as uniformly benign. Single-cell analysis reveals molecularly heterogeneous disease in which fibroblast transcriptional programs at biopsy are associated with divergent clinical trajectories: pathogenic myofibroblast modules associated with worse 52-week outcomes, and interferon-responsive and matrix-remodeling fibroblast programs with improvement. Because pathogenic and protective fibroblasts received overlapping immune-derived signals, our data are most consistent with the intrinsic transcriptional state of the fibroblast, rather than the inflammatory input it receives, shaping the stromal response. Baseline chronicity index captured this stromal heterogeneity and was the clinical parameter most associated with outcome. Together, these findings argue against a uniform watchful approach to Class II LN and identify baseline chronicity as a correlate warranting prospective evaluation.

## Data Sharing Statement

Single-cell RNA-seq data are available at Synapse (https://www.synapse.org/#!Synapse:syn26710600). Proteomics data will be deposited in publicly accessible, data-type-specific repositories and will be made publicly available as of the date of publication. All original code will be deposited in GitHub and will be made publicly available as of the date of publication. Any additional information required to reanalyze the data reported in this paper is available from the lead contact upon request.

## Author contributions

Conceptualization: K.V.R., J.S., J.P.B., B.H.R., A.F., M.P. Methodology: K.V.R., J.S., S.K., J.P.B., B.H.R., A.F., M.P., C.L. Investigation: J.S., K.V.R., S.K., J.P.B., B.H.R., A.F., M.P., C.L., L.L., R.M.C., H.S., T.T., T.M.E., B.J.M., M.W., D.M.F., M.G.A., J.M.T., R.F. Visualization: J.S., K.V.R., S.K., J.N.D. Funding acquisition: K.V.R., J.P.B., A.F., M.P. Project administration: K.V.R. Supervision: K.V.R., J.P.B., B.H.R. Writing – original draft: J.S., K.V.R., J.P.B., B.H.R. Writing – review and editing: all authors.

## Acknowledgements

We extend our gratitude to the NYU High Performance Compute (HPC) Center for their essential assistance with computational support and to the Clinical and Translational Science Institute (CTSI) for their crucial support in sample processing. We thank members of the Experimental Pathology Research Laboratory for their technical and imaging support. We appreciate Jennifer Seifert for her guidance within the AMP AIM consortium. We also thank clinical coordinators Katie Preisinger and Devyn Zaminski, as well as Benjamin Wainwright from the Buyon lab, for their support.

