## Supplemental figures for "Rethinking the Benign Nature of Class II Lupus Nephritis"

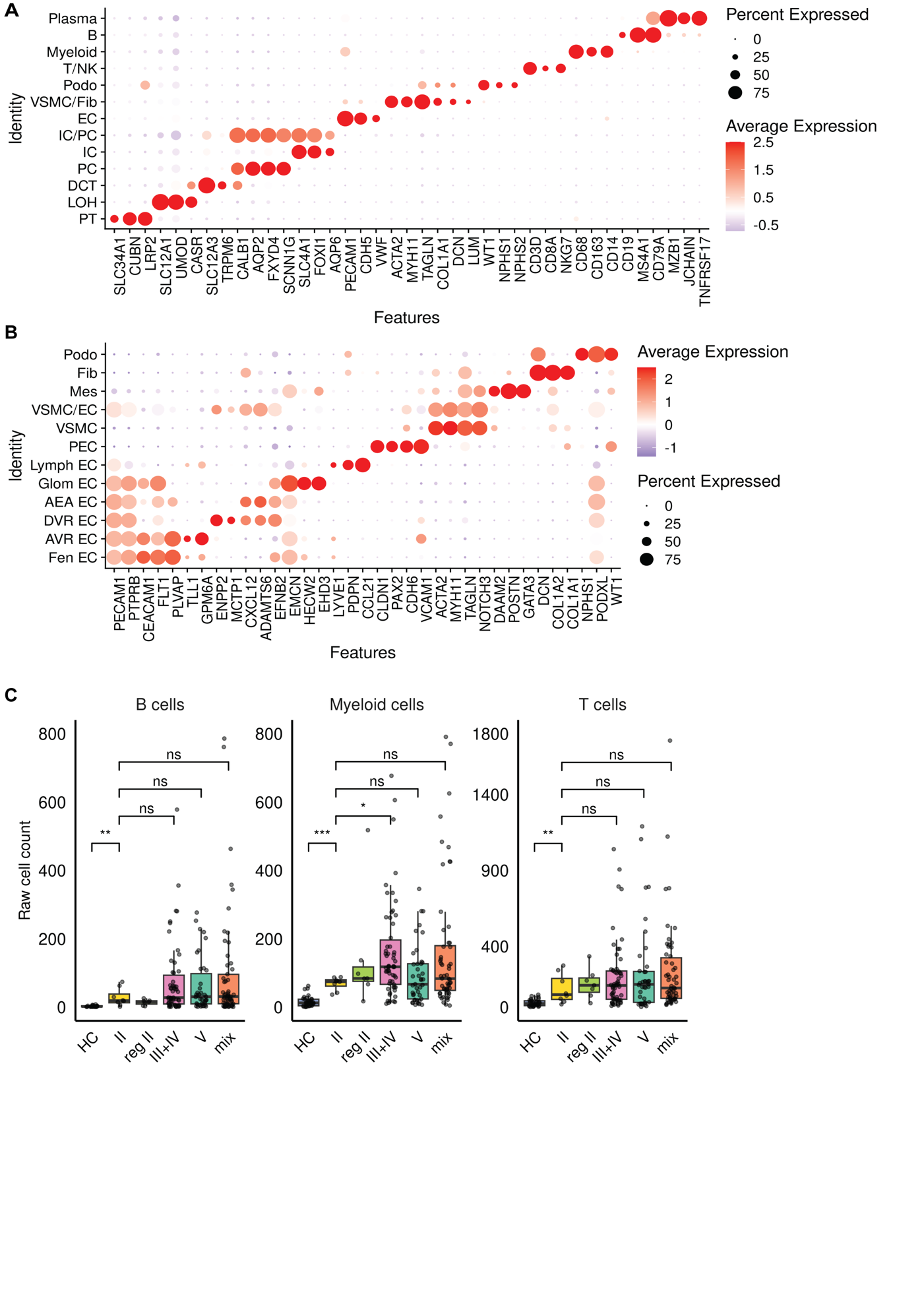

**Figure S1. Cell type markers and immune cell counts.** (**A**) Dotplot showing expression of canonical marker genes for cell types identified in the AMP dataset UMAP (Figure 1C, bottom). (**B**) Dotplot showing expression of canonical marker genes for cell types identified in the Class II dataset UMAP (Figure 1C, top). (**C**) Absolute immune cell counts (B cells, T cells, myeloid cells) per sample across disease states.

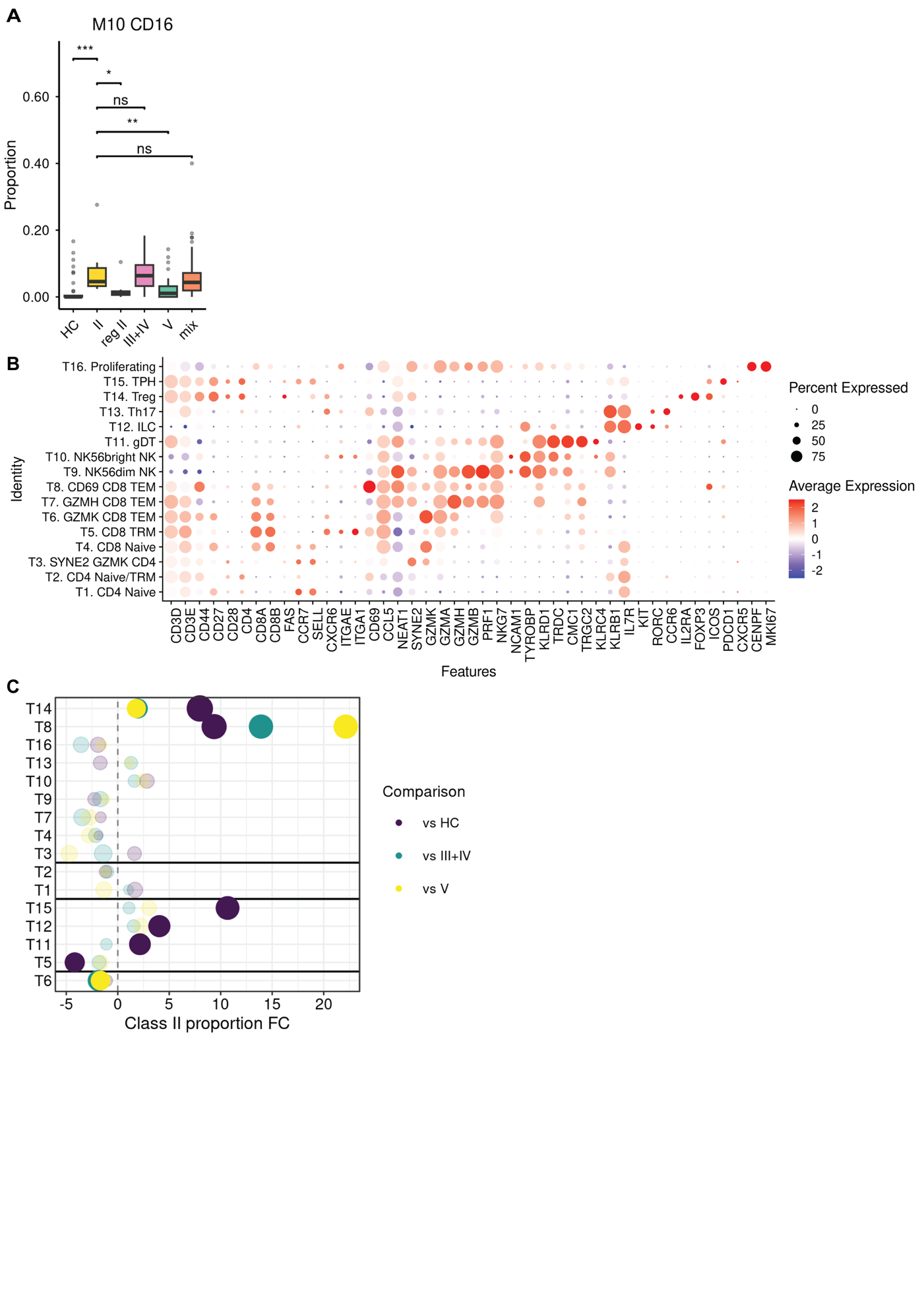
**Figure S2. Additional myeloid and T cell characterization.** (**A**) M10 CD16+ macrophage proportions across disease states. (**B**) Dotplot showing expression of canonical marker genes for T cell populations identified in Figure 2F. (**C**) T cell type proportional differences across disease states. Fold change (FC) comparing de novo Class II LN to healthy controls (HC), proliferative (Class III+IV), and membranous (Class V) LN. Dot size represents statistical significance (-log10 p-value) from Wilcoxon rank-sum tests, with larger dots indicating stronger significance. Colors distinguish between comparisons, while cell types are grouped based on whether their patterns are not significantly different from HC, shared with

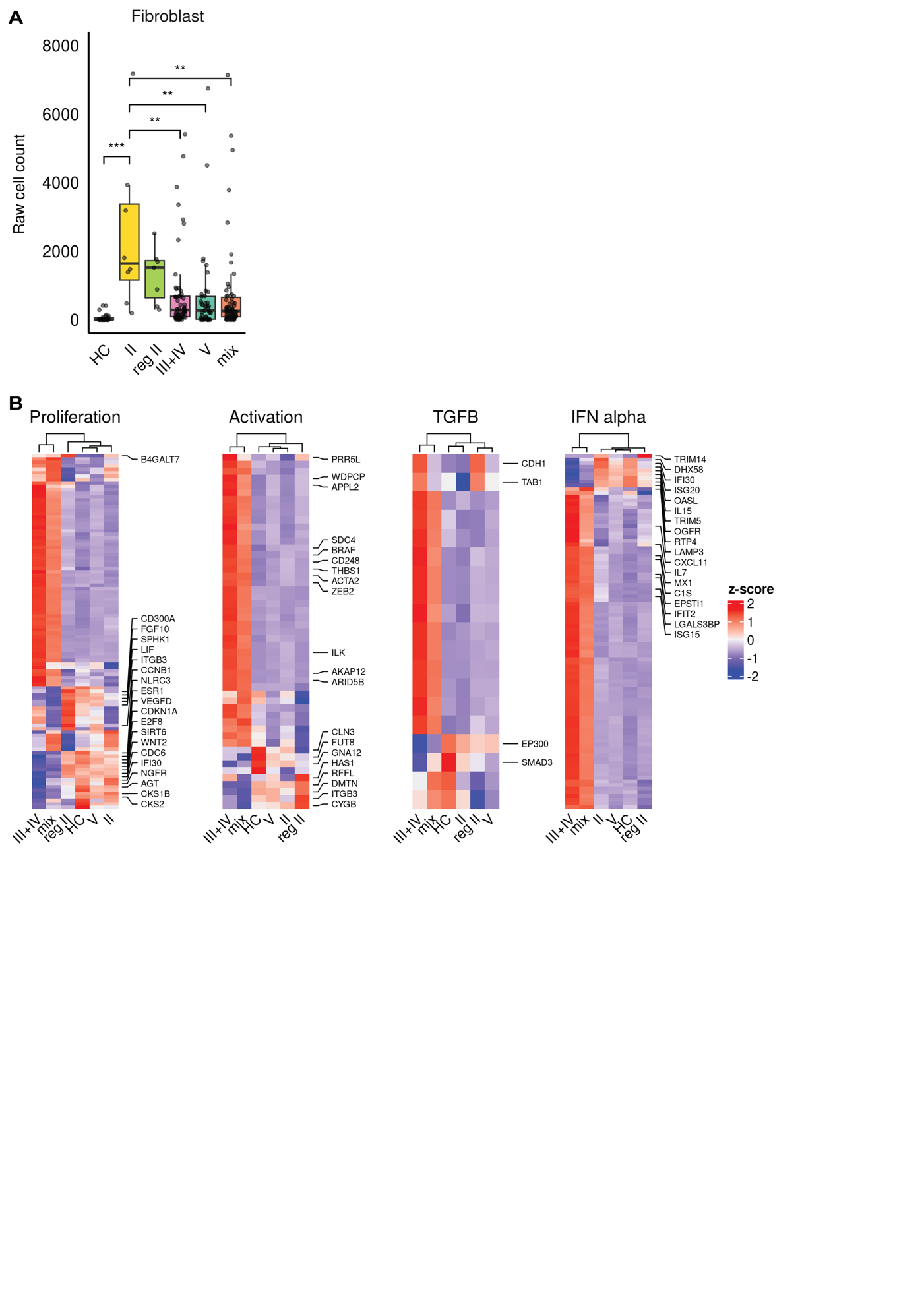

**Figure S3. Fibroblast and mesangial cell characterization.** (**A**) Absolute fibroblast cell counts per sample across disease states. (**B**) Per-class aggregated scaled expression in mesangial cells of proliferation/migration, activation, TGFβ signaling, and IFNα response gene signatures.

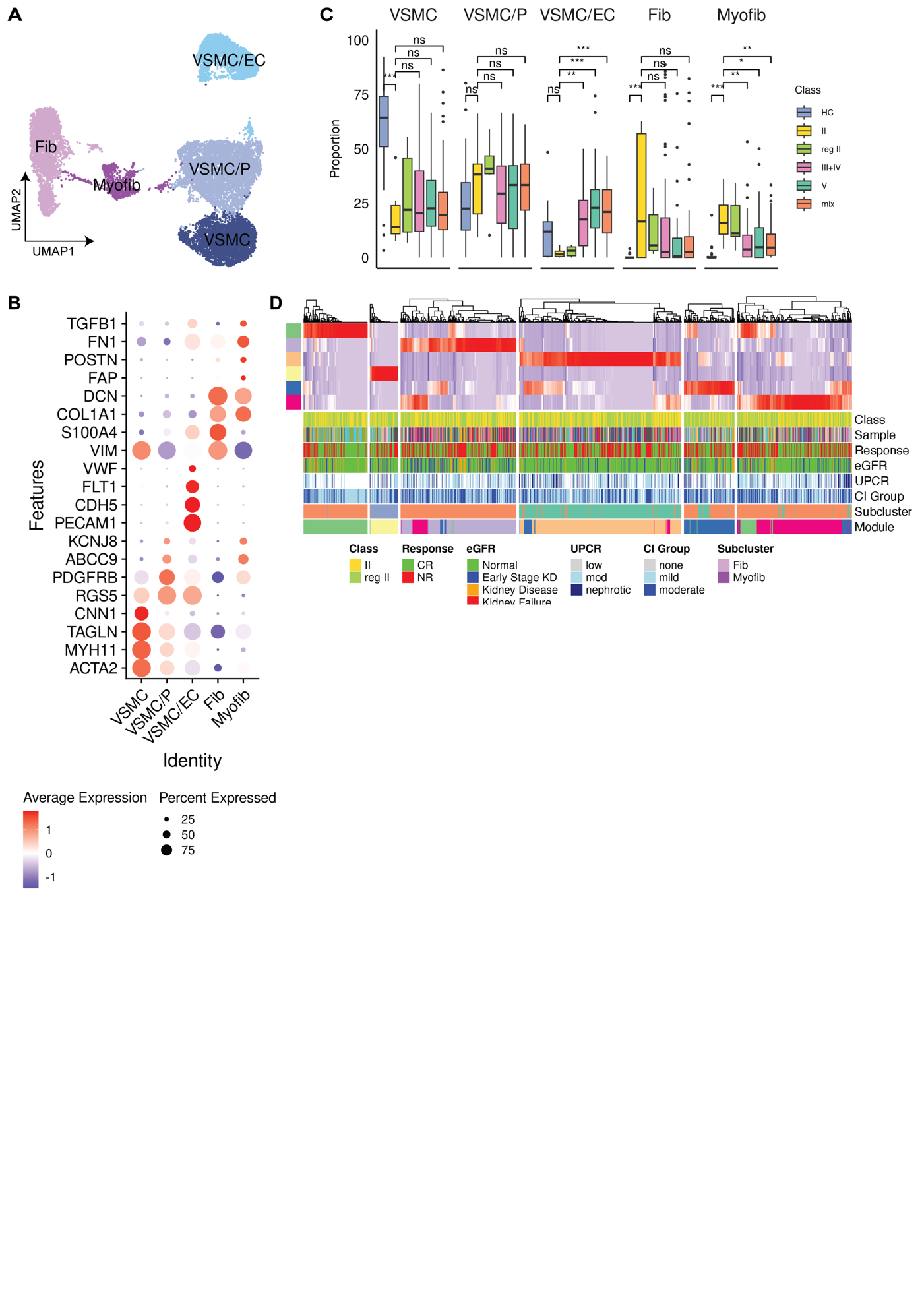

**Figure S4. Stromal cell characterization.** (**A**) UMAP visualization of fibroblasts, myofibroblasts, and VSMC populations. (**B**) Dotplot showing expression of defining markers for cell types in (A). (**C**) Per sample proportion of all cell types in (A) across disease states. (**D**) NMF gene module expression in VSMCs.
