## Supplemental methods for "Rethinking the Benign Nature of Class II Lupus Nephritis"

**Study approval and Patient recruitment**

**A total of 15 patients with Class II LN were included in this study.** Nine patients were enrolled in the AMP cohort; however, these patients were not included in prior AMP publications, which focused on membranous, proliferative, and mixed LN. Six additional patients with Class II LN were included in this study that met the AMP enrollment criteria: 1) age ≥ 18; 2) fulfillment of the revised American College of Rheumatology (1) or the Systemic Lupus Erythematosus International Cooperating Clinics (2) classification criteria for SLE; 3) a urine protein/creatinine ratio (UPCR) >0.5 at the time of biopsy. All patients provided written informed consent approved by their local institutional research board.

**Clinical data collection and patient classification**

As described in AMP (3), baseline demographics from a predetermined set of categories, including self-reported race (Asian, Black, White, Other)/ethnicity (Hispanic, non-Hispanic) and clinical characteristics were recorded at the time of biopsy. Laboratory tests and medications were documented at each visit (baseline, week 12, week 26, and week 52) and were performed at the participating sites. Based on established clinical guidelines, patients were categorized by eGFR (mL/min/1.73m²) into four groups: normal kidney function (≥90), early kidney disease (60-89), kidney disease (15-59), and kidney failure (<15); and by UPCR into three groups: low (<0.5), moderate (0.5-3.5), and nephrotic (>3.5).

**Kidney status assessment**

Kidney status was evaluated at 52-weeks post-biopsy using criteria adapted from the ACCESS Trial (4). Improvement was defined as: 1) UPCR < 0.5; 2) normal creatinine (≤ 1.3 mg/dL) or, if abnormal, ≤ 125% of baseline. Patients who did not improve by 52-weeks post biopsy could have remained stable (no changes in UPCR or serum creatinine), or could have worsened (increase in UPCR, serum creatinine, or both), but were grouped together as stable/worse. In accordance with the ACCESS trial protocol, microscopic review of the urine sediment was not included due to variability in assessment methods across sites and attribution challenges of hematuria in menstruating women.

**Tissue collection and processing**

As previously described (5), across all sites, ultrasound-guided or CT-guided kidney biopsies were performed by interventional radiologists or nephrologists using a 16-gauge to 18-gauge needle. In most cases, research tissue was obtained at the time of kidney biopsy as an additional core taken solely for research purposes, or in less than 15% by immediately obtaining a piece of a core with sufficient glomeruli that was not required solely for clinical diagnosis. All research tissue obtained was processed immediately and maintained in Cryostor for downstream tissue dissociation, sequencing and transcriptomic analysis.

**Pathological assessment**

Biopsies were reviewed by board certified pathologists at the institution where the biopsy was done and assigned histological classes according to the ISN/RPS classification (6,7). Although Class II is not typically graded using the NIH Activity and Chronicity Indices (AI and CI), given the complex previous biopsy history, these samples were also evaluated by one of the authors (BHR) for CI features of chronic damage such as Interstitial Fibrosis and Tubular Atrophy (IFTA) and Global Glomerulosclerosis (GSS). Samples were categorized into four groups based on chronicity scores: none (CI = 0), mild (CI = 1-3), moderate (CI = 4-7), or severe (CI ≥ 8) (**Fig. 1B**).

**Kidney tissue dissociation, single-cell isolation, DASH and sequencing**

Tissue samples were dissociated using a three-step enzymatic digestion protocol. First, samples were subjected to two sequential rounds of Liberase TL (Roche, 05401020001) treatment (0.25 mg/ml in PBS) for 15 minutes at 37°C. This was followed by a final digestion using 0.25% Trypsin for 10 minutes at 37°C. After each digestion step, the cell suspension was filtered through a 70 μm strainer into FBS and washed with PBS. The collected cells were pelleted by centrifugation at 300g for 5 minutes, followed by a wash step at 220g for 5 minutes in PBS containing 0.04% BSA. The final cell pellet was resuspended in PBS-BSA for cell counting and viability assessment. The final single-cell suspension was assessed for viability and cell count before proceeding to single-cell RNA sequencing.

After library preparation, mitochondrial transcript depletion was performed using Depletion of Abundant Sequences by Hybridization (DASH). Guide RNAs targeting 35 protein-coding mitochondrial transcripts and 26 mitochondrial rRNA sequences were synthesized by in vitro transcription from PCR-amplified DNA templates. Each 10X sequencing library (2 ng/μL) was treated with Cas9 nuclease (NEB, M0386T; 20 μM) and guide RNA pool (1.0 μM) in NEB3.1 buffer at 50-fold excess over library molecules. After pre-incubation at 25°C for 10 minutes, libraries were digested at 37°C for 2 hours to generate double-strand breaks. The reaction was terminated by RNase A (ThermoFisher) and Proteinase K (NEB, P8107S) treatment, followed by heat inactivation. Treated libraries were PCR-amplified for 6 cycles using KAPA HiFi HotStart ReadyMix (Kapa Biosystems, KK2602) and purified using AMPure XP beads (Beckman Coulter, A63881; 0.8X ratio). Cutting efficiency was assessed by qPCR using mitochondrial gene-specific primers before sequencing. Of note, MT-ND5 and MT-ND6 were intentionally left untargeted to serve as a proxy for overall mitochondrial transcript levels during RNA-seq quality control.

Single-cell RNA libraries were prepared using the 10x Genomics Chromium Single Cell 3' v3 chemistry (catalog # PN-1000268) following the manufacturer's protocol. Target cell recovery was 1500 cells per sample. Libraries were sequenced on the Illumina NovaSeq 6000 using an S2 flow cell with paired-end reads. Read 1 was 28 cycles (containing the cell barcode and UMI sequences), i7 index was 10 cycles, and Read 2 was 90 cycles (containing the transcript sequence). Sequencing parameters were set to achieve a minimum of 50,000 reads per cell.

**Quality control, filtering and pre-processing**

Raw sequencing reads were converted to count matrices with cell ranger’s (version 5.0.1) mkfastq and count functions, aligning to the GRCh38 human reference genome. For quality control of the single-cell RNA sequencing (scRNA-seq) data, each sample had cells removed with less than 500 unique genes detected and or less than 1000 UMI per cell. Additionally, cells with more than three percent of total reads from mitochondrial genes were removed, after accounting for DASH-targeted mitochondrial genes by specifically examining MT-ND5 and MT-ND6 expression.

scRNAseq data was processed using established quality control steps. First, doublets were identified and removed on a per sample basis using scDblFinder (v1.12.0, (8)). Two Class II samples (2726 and 2840) were excluded due to low quality metrics after mitochondrial and doublet filtering. Next, samples within each dataset (AMP and Class II) were merged separately and processed independently. Data normalization was performed using Seurat's SCTransform (v5.1.0, (9)) with regression of mitochondrial percentage. Principal component analysis was performed on the normalized data. Batch effect correction was implemented using Harmony integration (v1.2.0, (10)) on the first thirty principal components to account for variation from tissue procurement site, library preparation batch, and sample collection. Nearest neighbor graphs were constructed using the harmony-corrected principal components. Unsupervised clustering was performed using the Louvain algorithm and visualized using UMAP with thirty dimensions for each dataset.

**Broad cell type annotation**

Cell type identification was accomplished through a combination of known marker gene expression from the Kidney Precision Medicine Project (KPMP, (11)) and Azimuth (12) reference mapping, resulting in annotation of immune cell types and major kidney cell populations. For immune cell populations, T cells and NK cells were distinguished by expression of *CD3D*, *CD4*, and *NKG7*. Myeloid cells were characterized by expression of *CD14* and *CD68*, while B cells and plasma cells were identified through *CD79A* and *MS4A1* expression. Kidney parenchymal cells were classified based on segment-specific markers: proximal tubule cells expressed *LRP2* and *SLC34A1*, while loop of Henle cells were identified by *SLC12A1* and *CASR* expression, and the distal nephron segment was characterized by *CALB1*, *AQP2*, and *SLC4A1*. Endothelial cells were identified through *PECAM1* and *CDH5* expression, while fibroblasts and vascular smooth muscle cells (VSMCs) were distinguished by *DCN* and *ACTA2* respectively. Podocytes were identified through their expression of *NPHS1* and *WT1*.

**Identifying shared rare cell states**

After broad cell type identification, each major cell population was analyzed independently for finer cell type identification. For each cell population, the AMP dataset was subset, re-normalized, and subjected to dimensional reduction and clustering analysis using the same integration pipeline described above. To identify shared cell states between datasets, cells of the same broad identity in the Class II dataset were mapped to the AMP UMAP using Symphony (13). This approach enabled the identification of conserved cell states across both datasets while accounting for technical differences in sample processing. Lastly, canonical marker expression was assessed separately for AMP and Class II datasets to ensure shared cell type identity.

**Statistical and computational methods**

Statistical analysis of cell type composition across LN classes was performed using two complementary approaches: 1) Mann-Whitney U tests to compare median cell type abundances across LN classes within each cell type, and 2) linear regression models of the form *Proportion ~ ISN Class + Dataset*. The linear regression models tested for differences in cell type proportions between each LN class and controls while adjusting for dataset-specific effects. Models were fitted separately for each cell type using different minimum cell count thresholds to assess the robustness of the results.

To directly compare expression levels across datasets, batch effects were corrected using scGen (14), a deep learning-based generative model, trained for 100 epochs with early stopping. The trained model was then used to generate batch-corrected expression values while preserving biological variations associated with cell type identity. UMAP visualization of the corrected data demonstrated effective removal of batch-associated variation while maintaining cell type-specific expression patterns. Corrected counts were only used when directly comparing expression values in Fig 2D.

Differential gene expression analysis was performed on pseudobulk-aggregated single-cell data using the limma-voom framework. Raw counts were aggregated by sample within each cell type and filtered to retain genes with at least 5 counts in 30% of samples. Count data were normalized using the trimmed mean of M-values method and transformed to log2-counts per million using voom. A linear model incorporating processing batch as a covariate was fitted to the transformed data, followed by empirical Bayes moderation of standard errors. For scRNA-seq analysis, Fast Gene Set Enrichment Analysis (fgsea, v1.24.0) was selected as it performs optimally with the large number of genes and continuous expression values typical of transcriptomic data. Gene Ontology, Hallmark, and Reactome gene sets were used (15–17). Enrichment results were filtered based on statistical significance, magnitude of enrichment and biological relevance to kidney disease (complete results provided in Supplementary Tables 3 and 4). Cell-cell interaction networks were inferred using CellChat (v2.1.1, (18)) using default parameters.

**Non-negative Matrix Factorization, module annotation and scoring**

To identify distinct gene expression programs in Class II LN fibroblasts, we performed Non-negative Matrix Factorization (NMF) analysis. The input matrix was generated by applying SCTransform normalization and filtering to highly variable features (residual variance threshold of 1.3). Using the R package "NMF" we implemented the nsNMF method with random initialization, testing ranks 2-20 with 50 iterations each. The optimal rank was determined by evaluating the cophenetic correlation of both gene (basis) and sample (coefficient) consensus matrices. Genes were assigned to modules based on their highest value in the basis matrix and filtered using a threshold set at the median of normalized (0-1) module expression values. To facilitate comparison across LN classes while accounting for differences in sequencing depth, module gene lists were further filtered to include only genes identified as highly variable in the AMP dataset. Module activity scores were calculated using AUCell (19). Pathway enrichment analysis and manual geneset curation was performed to characterize the biological functions of each module.

**Immunofluorescence and imaging**

Multiplex immunofluorescence staining was performed on a Leica BondRx autostainer with Akoya Biosciences® Opal™ reagents, according to the manufacturers’ instructions. 5 um thick sections were treated with hydrogen peroxide to inhibit endogenous peroxidases followed by an antigen retrieval step with ER2 (pH9, Leica 9AR9640) for 20 minutes at 100°C. Slides were incubated with the anti-S100A4 primary antibody (CST, 13018S,1:1000 dilution) followed by the anti-mouse + rabbit secondary polymer (Akoya, ARH1001EA). Slides then underwent HRP-mediated tyramide signal amplification with the 620 Opal® fluorophore (Akoya, FP1495001, 1:150 dilution). The primary and secondary antibodies were subsequently removed with a heat retrieval step, leaving the Opal fluorophore covalently linked to the antigen. This sequence was repeated with the Ki67 primary antibody and Akoya secondary polymer with the 690 Opal fluorophore (Akoya FP1497001, 1:150 dilution). Sections were counterstained with spectral DAPI (Akoya Biosciences, FP1490) and mounted with ProLong Gold Antifade (ThermoFisher Scientific, P36935). Semi-automated image acquisition was performed on a Akoya Vectra Polaris (PhenoImagerHT) multispectral imaging system at 20X magnification using PhenoImagerHT 2.0 software in conjunction with Phenochart 2.0 and InForm 3.0 to generate unmixed whole slide qptiff scans. The image files were uploaded to the NYUGSoM’s OMERO Plus image data management system (Glencoe Software).

**Urine collection, processing and analysis**

Urine was collected from AMP patients 52-weeks post biopsy as previously described (20). Of the fifteen Class II samples, only seven were matched to scRNAseq of kidney tissue, while the other eight were from Class II patients not included in the scRNAseq analysis. Briefly, protein abundances were quantified using the Kiloplex Quantibody protein array platform (RayBiotech, (21)), capturing over 1,200 proteins. Protein abundances were log-transformed after adding a small constant (10% of the minimum measured abundance) to eliminate zero values, enabling normalization of the data distribution. Differential protein expression between lupus nephritis classes (proliferative, mixed, membranous, and class II) and healthy controls was assessed using Wilcoxon rank-sum tests. P-values were adjusted for multiple comparisons using the Benjamini-Hochberg method. Different enrichment tools were used to accommodate the distinct characteristics of scRNAseq and proteomics data. For proteomic analysis, enrichR (v3.2, (22)) was utilized as it is better suited for the smaller, discrete sets of proteins typically identified in proteomic experiments. Comparisons between LN classes and healthy controls were performed using Mann-Whitney U tests.
